# Experimental evolution for niche breadth in bacteriophage T4 highlights the importance of structural genes

**DOI:** 10.1101/669770

**Authors:** Jenny Y. Pham, C. Brandon Ogbunugafor, Alex N. Nguyen Ba, Daniel L. Hartl

## Abstract

Ecologists have long studied the evolution of niche breadth, including how variability in environments can drive the evolution of specialism and generalism. This concept is of particular interest in viruses, where niche-breadth evolution may explain viral disease emergence, or underlie the potential for therapeutic measures like phage therapy. Despite the significance and potential applications of virus-host interactions, the genetic determinants of niche-breadth evolution remain unexplored in many bacteriophage. In this study, we present the results of an evolution experiment with a model bacteriophage system, *Escherichia virus T4*, in several host environments: exposure to *E. coli* C, exposure to *E. coli* K-12, and exposure to both *E. coli* C and *E. coli* K-12. This experimental framework allowed us to investigate the phenotypic and molecular manifestations of niche-breadth evolution. First, we show that selection on different hosts led to measurable changes in phage productivity in all experimental populations. Second, whole—genome sequencing of experimental populations revealed signatures of selection. Finally, clear and consistent patterns emerged across the host environments, especially the presence of new mutations in phage structural genes. A comparison of mutations found across functional gene categories revealed that structural genes acquired significantly more mutations than other categories. Our findings suggest that structural genes—those that provide morphological and biophysical integrity to a virus—are central determinants in bacteriophage niche breadth.

## Introduction

Niche breadth reflects the range of resources or habitats used by a given species or population (Sexton, Montiel, Shay, Stephens, & Slatyer, 2017). It can be defined by a continuum with two extremes: specialists, which have maximized fitness on one resource, and generalists, which display similar fitness over a broad range of resources (Sexton et al., 2017). Ecological theory offers the prediction that populations persisting in a stable environment will evolve a specialist strategy, where performance in that single environment is the target of selection (Wilson & Yoshimura, 1994). In contrast, temporally variable environments may favor the evolution of generalists that are able to tolerate the range of encountered environments (Levins, 1968). Theory also proposes a tradeoff between niche breath and fitness on a particular resource; specialists suffer reduced performance on alternate resources whereas generalists cannot exploit a particular resource as efficiently as the specialist (Futuyma & Moreno, 1988; Lynch & Gabriel, 1987; Palaima, 2007). Two proposed genetic mechanisms responsible for tradeoffs include antagonistic pleiotropy (i.e., mutations are beneficial in one environment and detrimental in others) and mutation accumulation (i.e., mutations are neutral in the environment in which they arose but are detrimental in others) (Siobain Duffy, Turner, & Burch, 2006; Kawecki, 1994; MacLean, Bell, & Rainey, 2004).

Experimental evolution has served an important role in the study of niche breadth in microbes (Kassen, 2002). Many of these studies have focused on virus systems and experimental exposure to different (or new) hosts, as the type, range, and availability of hosts present in the environment is an important source of selection pressure (Cooper & Scott, 2001; Crill, Wichman, & Bull, 2000; S. Duffy, Burch, & Turner, 2007; Kutnjak, Elena, & Ravnikar, 2017; Morley, Mendiola, & Turner, 2015; Ogbunugafor et al. 2013, Novella et al., 1995). This and other experimental work in this realm has provided varying levels of support for the tradeoff hypothesis, indicating that there are still gaps in our knowledge regarding how viruses respond to changes in their environment (Bedhomme, Lafforgue, & Elena, 2012; Novella, Hershey, Escarmis, Domingo, & Holland, 1999; Turner & Elena, 2000). In particular, there have been relatively few rigorous treatments of the evolutionary genomics of niche-breadth expansion, where the genetic determinants are fully resolved with functional inferences drawn between mutations and phenotypes. Such studies continue to have practical relevance for questions about the phenotypic manifestations of niche-breadth evolution, whether tradeoffs arise as a consequence of adaptation, and about the molecular signatures of such evolution.

In this study, we sought to answer these questions by characterizing the phenotypic and molecular changes associated with niche-breadth evolution in a model bacteriophage. We employ *Escherichia virus T4* as a system for experimental evolution. T4 is of particular interest because it is among the most well-studied and fully characterized viruses. T4 is also surprisingly complex, with a genome ~170 kb in size and a gene density four times greater than that of herpes viruses and twice that *Escherichia coli*. Of its 300 genes, approximately 160 have been functionally characterized (Miller et al., 2003), which provides the opportunity to resolve molecular mechanisms responsible for niche-breadth evolution. We evolved T4 for 50 generations in one of three host environments: 1) *E. coli* C, 2) *E. coli* K-12, and 3) daily alternation of *E. coli* C and *E. coli* K-12 (Fig. 1).

**Figure 1:**
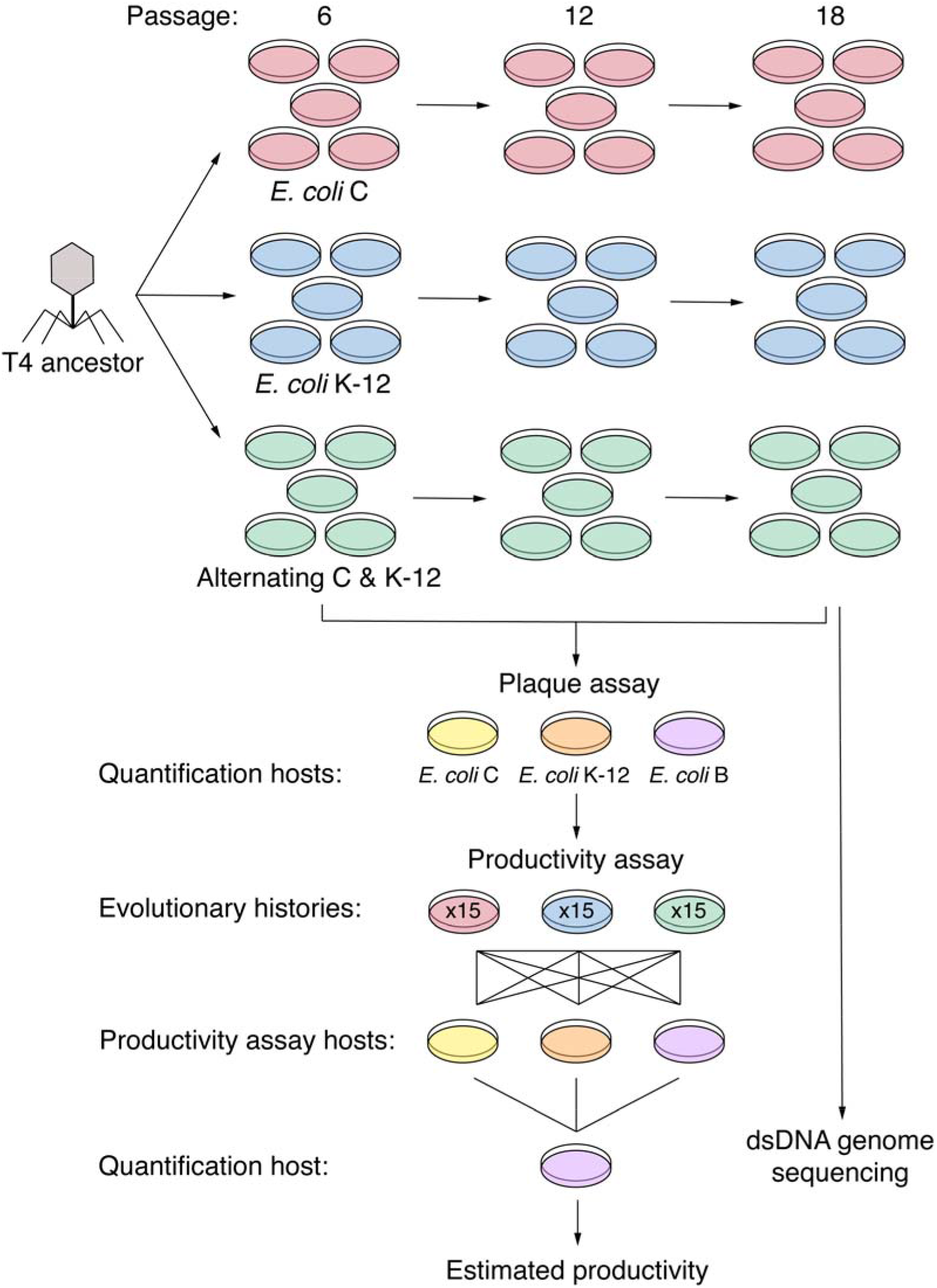
Experimental evolution schematic. 15 populations were seeded with a previously isolated T4 ancestral clone and split into three evolutionary histories with different host environments: five populations were exposed to *E. coli* C, another five were exposed to *E. coli* K-12, and the last five were exposed to *E. coli* C and K-12 in daily alternation. Serial passaging occurred for 20 days, which is equivalent to approximately 50 generations. Plaque assays on the original host *E. coli* B and the selection host *E. coli* C and/or K-12 were performed for quantification of evolved phage samples on passages 6, 12, and 18. To measure productivity, assays on *E. coli* C, K-12, and B were performed for the same evolved samples. Following the productivity assay, samples were quantified using *E. coli* B, which generated the final estimate of productivity (log_10_ titer [pfu/mL]). Sequencing was performed on the complete genomes of the T4 ancestor and 15 evolved populations at passage 18.

Selection on either *E. coli* C or *E. coli* K-12 mimics a constant environment, which is predicted to drive the evolution of specialists; whereas selection on the alternating hosts mimics a temporally variable environment, which is predicted to drive the evolution of generalists (Turner & Elena, 2000). Our results reveal the complexity of niche-breadth evolution, with some populations demonstrating properties of a tradeoff, others less so. The genomic data reflected patterns across evolutionary histories: new mutations were overrepresented in genes that encode structural virion proteins. Notably, this pattern implies that structural genes—and in particular, those that function in host recognition, infection, and stability—are important in niche-breadth evolution, regardless of conditions that promoted a particular ecological strategy (specialism or generalism). We discuss these findings in detail, and reflect on their implications for general viral ecology, and for the various arenas where bacteriophage niche breadth has practical utility—in disease emergence, public health surveillance, and efforts to engineer bacteriophage for therapeutic purposes.

## Materials and Methods

### T4 and bacterial strains

This study used *Escherichia virus T4* (American Type Culture Collection [ATCC] #11303-B4) and three wild-type bacterial hosts: *E. coli* B (ATTC #11303), *E. coli* C (Coli Genetic Stock Center 3121), and *E. coli* K-12 (Coli Genetic Stock Center 4401). T4 infection of *E. coli* K-12 has been well documented (Yu & Mizushima, 1982), but not *E. coli* C, which is a strain normally used for the propagation of *Escherichia virus phiX174* (Wichman, Millstein, & Bull, 2005). *E. coli* B is the strain that has been historically used for the propagation of T4 (Demerec & Fano, 1944) and currently recommended by the ATCC for T4 propagation. Bacteria were stored as 25% glycerol stocks at −80°C; isolated bacterial colonies were obtained by streaking stocks onto Luria-Bertani (LB) agar petri dishes (BD, VWR International). All assays and serial passaging utilized liquid bacterial cultures which were prepared daily; single colonies were inoculated into glass culture tubes containing 5 ml of LB broth and incubated overnight at approximately 180 RPM (C1 Platform Shaker, New Brunswick Scientific) and 37°C. The T4 ancestral clone used to initiate experimentally evolved lines was isolated by plaque purification on LB agar petri dishes containing overlays comprised of 10 μl serially diluted phage lysate, 30 μl (~10^8^ colony forming units [CFU]/ml) bacterial host culture, 270 μl LB broth, and 4 ml LB soft agar (7.5 g/L agar). Following overnight incubation at 37°C, a single well-formed plaque was isolated, saturated with 500 μl LB broth, treated with 4% chloroform (VWR Life Science), and stored at 4°C.

### Experimental evolution of T4

To initiate the experimental evolution, the T4 ancestral clone was used to seed 15 populations. Three sets of five parallel populations were evolved in one of three host environments: 1) *E. coli* C, 2) *E. coli* K-12, and 3) a daily alternating pattern of *E. coli* C and *E. coli* K-12 (Fig. 1). The initial passage was performed in LB agar petri dishes containing overlays comprised of the T4 ancestral clone, appropriate bacterial host, 270 μl LB broth, and 4 ml LB soft agar, with initial infection occurring at MOI = 0.001 (volumes for T4 and bacteria were variable due to differences in host growth). Following overnight incubation at 37°C, phage lysates were collected, treated with 4% chloroform, centrifuged at 7500 rpm and 4°C for 20 minutes (J6-MI, Beckman Coulter), and sterilized using 0.2 μm syringe filters (Pall Life Sciences). 500 μl of each phage lysate was saved for storage at 4°C. These steps ensured that all bacteria were removed from phage lysates prior to usage for subsequent passages. Passages following the initial infection were plated as overlays on LB agar petri dishes, which contained 10 μl of phage lysate, 30 μl bacterial host culture, 270 μl LB broth, and 4 ml of LB soft agar, followed by overnight incubation at 37°C. Lysates generated over the course of the experimental evolution were collected, purified, and stored in the same manner as the initial lysate. In all passages, phage populations were propagated with hosts derived from newly prepared overnight bacterial cultures (no possibility for coevolution). To prevent cross-contamination between evolutionary histories, each set of parallel populations was confined to separate laboratory rooms and lysate supernatants were collected and purified using only materials (i.e., micropipettes and plastics) contained to the same room. Serial passaging was performed for 20 days, which is assumed to be equivalent to ~50 generations of T4 growth (we elaborate on generation time below).

### Determination of generations

Due to exponential growth via binary fission, the number of bacterial generations (*G*) is calculated by the equation below, with *B*_0_ representing the initial number of bacteria and *B*_*F*_ representing the number of bacteria at the end of a time interval (Lenski, Rose, Simpson, & Tadler, 1991; Monod, 1949):

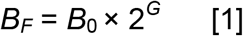

Bacteriophage also grow exponentially, with the number of generations being determined by burst size (the number of phage progeny produced per infected bacterium) rather than binary fission. The bacterial equation can then be modified to suit phage, with *V*_0_ representing the initial number of phage, *V*_*F*_ representing the number of phage at the end of a time interval, and *R* representing burst size (Abedon et al., 2001; Miralles, Moya, & Elena, 2000):

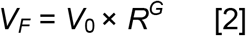

The burst size for T4 on *E. coli* B has been previously determined to be approximately 110 PFU/ml (Hadas, Einav, Fishov, & Zaritsky, 1997) and a typical overnight infection of T4 in *E. coli* B results in 100,000× growth (based on experimental data from Fig. 2), hence the number of generations is ~2.5. Though this calculation can provide a general impression of the number of T4 generations which occurred during experimental evolution, it is an imperfect estimate. A more accurate estimate would require knowledge of the burst size on the selected host (which may be different than the burst size on *E. coli* B) and the number of phage added and produced during serial passaging infections (which may change from one passage to another).

**Figure 2:**
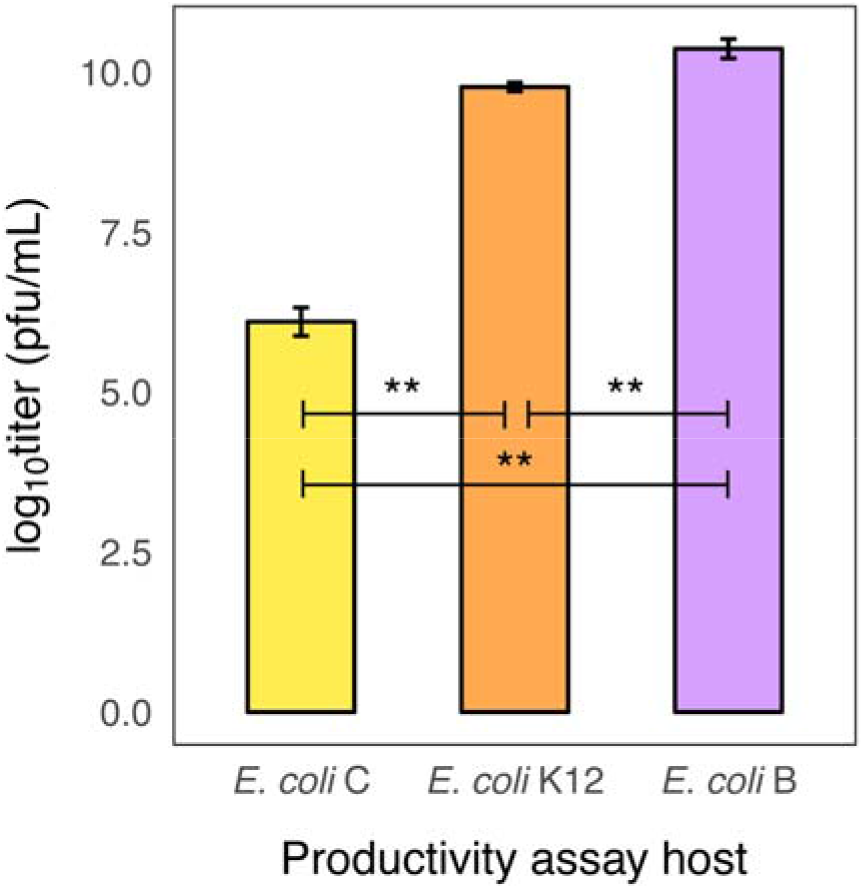
Productivity of the T4 ancestral clone. Productivity of the T4 ancestral clone was measured on the experimental evolution hosts, *E. coli* C and K-12, and on the original host, *E. coli* B. Each bar represents mean productivity (log_10_ titer [pfu/mL]) measured over three replicate assays and error bars indicate 95% confidence limits. Analysis using one-way ANOVA followed by Tukey’s HSD test indicated that all comparisons of mean titer were significant (*P* < 0.01).

### Phage quantification

Quantification was accomplished using plaque assays in which 4 μl of 10-fold serial dilutions of phage lysates were spotted on LB agar petri dishes containing overlays comprised of 30 μl bacterial host culture, 270 μl LB broth, and 4 ml LB soft agar. Petri dishes were incubated overnight at 37°C, with plaque visualization and counting occurring the following day. Quantified phage samples were expressed as PFU/ml; each plaque was assumed to have originated from a single infecting phage particle, thus one plaque was equivalent to one PFU. Throughout the experiment, *E. coli* B was used for quantification because it is a consistent and highly permissive host. Following the experimental evolution, three replicate plaque assays (*n* = 3) were performed on the 15 populations at passage 6, 12, and 18, using both the selection host/hosts and the original host, to confirm that *E. coli* B remained a more sensitive host. This yielded 315 data points ([5 populations × 2 single host histories × 3 time points × 2 quantification hosts × 3 replications] + [5 populations × 1 alternating host history × 3 time points × 3 quantification hosts × 3 replications]). Replicate titers were log_10_-transformed and compared for differences due to quantification host using unpaired *t*-tests; each comparison was comprised of 30 measurements because titers were pooled by evolutionary history. In total, 12 comparisons were made using GraphPad Prism v. 7.0b (GraphPad Software, La Jolla California USA).

### Productivity assays

Productivity assays were performed to measure phage titers produced on three hosts: *E. coli* C, *E. coli* K-12, and *E. coli* B. Following the experimental evolution, assays were performed on the 15 populations at passage 6, 12, and 18. Productivity assays were standardized by infection of phage samples and bacteria at MOI = 0.001 and measured total number of progeny produced. Each assay was performed on LB agar petri dishes containing overlays comprised of 10 μl diluted phage lysate, 1 μl diluted bacterial host, 270 μl LB broth, and 4 ml LB soft agar. Following overnight incubation at 37°C, phage lysates were collected and purified using the same methods as those applied to the serial passages. Lastly, phage lysates were quantified for titer using plaque assays (as described above) and *E. coli* B as the quantification host.

In total, three replicate productivity assays (*n* = 3) were performed, which resulted in 405 titer measurements (5 populations × 3 evolutionary histories × 3 time points × 3 assay hosts × 3 replications). Titer measurements used in the linear mixed effects model were log_10_-transformed to improve normality. The model was generated using the lmer function of the lme4 R package (Bates, Mächler, Bolker, & Walker, 2015). The initial model included replication as an additional random factor, however this parameter was dropped because the variance estimate was extremely small (*variance* = 0.00142) when compared to the variance of the residual error (*variance* = 0.11195). To confirm goodness of fit and ensure that model assumptions were met, model residuals were visually inspected (Zuur, Ieno, & Elphick, 2010). Plotted residuals were normally distributed and revealed no concerning patterns of heterogeneity in variances. The analysis of variance table was calculated using the Anova function of the car R package, which performs Wald chi-square tests for linear mixed effects models (Fox & Weisberg, 2011). Follow-up analysis was performed using the contrast function of the lsmeans R package to obtain pairwise comparisons among LS means (Lenth, 2016). *P* values generated from this analysis were adjusted using the Holm method, which is designed to give strong control of the family-wise error rate while retaining more power than the Bonferroni correction. The results of this analysis are available in Supplementary Table 1a and 1b.

Additional productivity assays (*n* = 3) were performed for the T4 ancestor in order to gauge baseline titer on the three assay hosts. These data were excluded from the linear model and analyzed using one-way ANOVA followed by Tukey’s HSD test. Titer measurements of the ancestor were necessarily excluded from the linear model because by definition, the ancestor did not have an evolutionary history nor was it serially passaged. All above analyses were performed in R v. 3.3.3 (R Development Core Team, 2017).

### Phage DNA extraction

Phage lysates were newly generated from the original evolved phage samples for library preparation. This step was performed for two reasons: 1) to preserve the original 500 μl generated directly from the serial passaging and 2) phage DNA extraction protocols generally require at least 1 ml of high titer (> 10^9^) phage lysate to generate high yield, clean DNA. Lysates were created from evolved samples at passage 18, the final time point that was assayed for productivity (Fig. 1). These were plated as overlays on LB agar petri dishes, which contained 10 μl of evolved phage sample, 30 μl bacterial host culture, 270 μl LB broth, and 4 ml of LB soft agar. Following overnight incubation at 37°C, lysates were collected and purified in the same manner as with the serial passaging.

1 ml of each lysate was first treated with 12.5 μl 1M MgCl_2_ (VWR Life Science), 0.4 μl DNAse 1 (2000 U/ml) (New England Biolabs), and 10 μl RNAse A (10 mg/ml) (ThermoFisher Scientific). Following brief vortexing and room temperature (RT) incubation for 30 minutes, the following reagents were added: 40 μl of 0.5 M EDTA (Corning), 2.5 μl Proteinase K (20 mg/mlL) (ThermoFisher Scientific), and 25 μl 10% SDS (VWR Life Science). The mixture was vortexed vigorously and incubated at 55 °C for 60 minutes, with additional mixing occurring twice at 20 minute intervals. Phage DNA was extracted using an equal amount of phenol:chloroform:isoamyl alcohol (25:24:1) (VWR Life Science) to lysate. Following inversion mixing and centrifugation at RT for 5 minutes at 13K, the aqueous layer was removed and two further extractions were performed in the same manner. Phage DNA was precipitated by addition of 1 ml 90% ethanol (Pharmco) and 50 μl 3M sodium acetate (Corning). Following incubation on ice for 5 minutes, DNA was gently mixed and centrifuged at room temperature for 10 minutes at 13K. The DNA pellet was washed with 500 μl 70% ethanol and centrifuged at RT for 10 minutes at 13K. The ethanol was decanted and the pellet was air dried for ~20 minutes. DNA was reconstituted in 50 μl TE buffer (Invitrogen) and checked for purity and concentration with Nanodrop 2000 (Thermo Scientific).

### Library preparation and sequencing

Library preparation was executed according to previously published protocols (Baym et al., 2015). Minor protocol modifications were made and are thus noted. In module 1, DNA was standardized with Qubit dsDNA HS Assay Kit (ThermoFisher Scientific). In module 4, DNA size selection was performed twice (rather than once). First, 0.55× magnetic beads were added to genomic DNA to remove large (> 600 bp) fragments. Following incubation at RT for 5 minutes, the tubes were placed on a magnetic stand and the supernatant was removed and transferred to new PCR strip tubes. In the second size selection, 0.3× magnetic beads were added to the supernatant to bind DNA fragments of desired length (~250 – 600 bp). Following incubation at RT for 5 minutes, tubes were placed on a magnetic stand and the supernatant was removed and discarded. The beads were washed once with 200 μl 80% ethanol, allowed to air dry for ~20 minutes, and resuspended in 30 μl TE buffer. In module 4, DNA was quantified using Qubit dsDNA HS Assay Kit; libraries were pooled at 5 ng per sample and fragment size was assessed with Agilent 4200 TapeStation System. The final pooled library sample was sent to Bauer Core Facility (Harvard University, Cambridge, MA) for QPCR quality control and sequencing using Illumina NextSeq 500 Mid Cycle (2 x 150 bp).

### Analysis of sequence data

Demultiplexed reads were trimmed for Nextera adapter sequences using Trimmomatic with default settings (Bolger, Lohse, & Usadel, 2014). The reads were aligned to the T4 reference genome (RefSeq accession no. GCF_000836945.1) using Breseq v. 0.31.1 (Deatherage & Barrick, 2014). For populations, the polymorphism mode was enabled, and no junction predictions were made. Typical coverage depth was over 1000× for most positions in the genome with over 95% of the reads mapping. The ancestral clone sample was replicated in the library preparation stage to account for sequencing errors and potential variation between the reference genome and the ancestral clone. Following sequencing, mutations were called separately for the two ancestral clone samples; these replicates revealed complete agreement between the called mutations. The ancestral mutations were subsequently filtered from all evolved populations in order to account for mutations that were present in the T4 ancestor but absent in the reference genome. The default Breseq configuration requires a mutation to be found in at least 5% of the reads, hence only SNPs present in the populations at 5% or greater were considered. All polymorphisms and their frequencies are available in Supplementary Table 2a.

In the present analysis, sequencing error bias presented barriers for the determination of polymorphisms. To remedy this, mutations with consistent statistical evidence of strand bias were removed in circumstances where the mutation would not be called when considering information from one strand versus the other. Specifically, mutations were filtered due to strand bias in cases where there existed at least 5% difference between the forward and reverse strand (preventing cases where a mutation is found in 4.9% on one strand, and 5.1% on the other one, but filtering out cases where a mutation is found in 1% in one strand, but 6% on the other). xNPs similarly presented issues with the NGS analysis pipeline, as it did not recognize sequential mutations as one mutational event, but rather multiple events. This issue manifests in the context of calling amino acid changes, i.e., when two adjacent mutations occur within one codon, two different amino acid changes are detected for each mutation, rather than one amino acid change which incorporates both nucleotide substitutions. In these cases, reads were visualized to ensure all putative xNPs occurred within the same read (as opposed to a situation where mutations occurred separately in different reads) to validate the presence of xNPs. The amino acid calls were subsequently manually modified to reflect the true amino acid change. A similar issue occurred with indels; each indel was reported as multiple mutational events rather than one. These situations were resolved in the same manner as xNPs. A final issue occurred in which a fixed ancestral mutation was detected as a new polymorphism, when it decreased in frequency due to a new mutation. This new mutation was not detected because the nucleotide substitution was coincidentally the same base as that of the reference sequence used for alignment. This error was confirmed and manually corrected by visualizing reads containing the relevant position. A list of corrected positions is available in Supplementary Table 2b.

### Analysis of mutation distribution

The number of nucleotide sites occupied by each functional category was determined using previous gene designations (Miller et al., 2003) and the T4 genomic sequence associated with that study (Genbank accession no. AF158101.6). From this, the mutation rate (number of mutations per nucleotide site) of each functional gene category was calculated for the 15 evolved populations. All polymorphisms detected at or above 5% were considered for this analysis. A SRH non-parametric two-way ANOVA (Sokal & Rohlf, 2011) was performed to test the influence of factors evolutionary history, functional gene category, and their interactions on rate of mutation. This analysis was first performed for all mutations, and then repeated separately for synonymous and nonsynonymous mutations. The analyses were executed using the scheirerRayHare function of the rcompanion R package (Mangiafico, 2018). Follow up analysis of pairwise comparisons was performed using Dunn’s test, which was executed using the dunnTest function of the FSA R package (Ogle, 2018). *P* values generated from the pairwise comparisons were adjusted using the Holm method. All above analyses were performed in R v. 3.3.3 (R Development Core Team, 2017).

## Results

### The baseline productivity of the T4 ancestral clone differs according to host strain

Before discussing the results corresponding to the state of phage populations post-experimental evolution, we first examined the baseline productivity of bacteriophage T4 across the various hosts (*E. coli* C, *E. coli* K-12, and *E. coli* B) used in the experimental evolution. Phage productivity was chosen as an estimate for gauging performance on a particular host because it represents the number of infectious progeny produced (per infected bacterium or per parent virion) from an infection (Abedon, Herschler, & Stopar, 2001; Kerr, Neuhauser, Bohannan, & Dean, 2006). The ancestral clone displayed differing levels of productivity on each host strain, which was expressed as mean titer (log_10_ plaque forming units [PFU]/ml). Productivity was predictably highest on the ancestral host *E. coli* B (10.37 log_10_ PFU with 0.14 standard deviation [SD]), slightly lower on *E. coli* K-12 (9.77 log_10_ PFU, 0.06 SD), and lowest on *E. coli* C (6.10 log_10_ PFU, 0.20 SD) (Fig. 2). Analysis of these data using one-way ANOVA followed by Tukey’s Honest Significant Differences (HSD) test indicated that productivity differences were significant (*P* < 0.01). Notably, T4 productivity was slightly different but high on *E. coli* K-12 and B, while productivity on *E. coli* C was several orders of magnitude (on the order of 10^4^) lower than the others.

### After evolution, *E. coli* B remained the most sensitive phage quantification host

Following experimental evolution, samples required quantification of phage titer prior to being assayed for productivity. *E. coli* B is commonly used for T4 quantification because it is a consistent and highly permissive host. We anticipated a potential issue because selection on other hosts could lead to correlated antagonistic changes (*i.e.*, reduced infectivity) on the original host. Plaque assays were performed on the 15 populations at three time points (passage 6, 12, and 18) to investigate whether *E. coli* B remained a more sensitive quantification host than the selected host(s) (*E. coli* C and/or *E. coli* K-12). Results from these plaque assays indicated that phage titer measurements, expressed as log_10_ PFU/mL, on *E. coli* B were either equivalent or greater than those on the selected host(s) (Fig. 3). Specifically, the grand-mean titer for *E. coli* C evolved populations was higher on *E. coli* B than that on *E. coli* C for all time points, indicating that *E. coli* B was a more sensitive quantification host. The grand-mean titer for *E. coli* K-12 evolved populations was equivalent whether quantified on *E. coli* B or K-12 for all time points, indicating that these two strains were equally sensitive as quantification hosts. The grand-mean titer for evolved populations with alternating hosts recapitulated the results of the single-host evolved populations. For all time points, *E. coli* B provided equivalent titer measurements when compared to *E. coli* K-12 and more sensitive measurements when compared to *E. coli* C. Furthermore, a series of *t*-tests conducted on titer measurements pooled by evolutionary history indicated that all comparisons involving *E. coli* C and *E. coli* B as quantification hosts yielded significant differences (with the higher titer estimate occurring on *E. coli* B) whereas all comparisons involving *E. coli* K-12 and *E. coli* B yielded no significant differences (Table 1). These results confirm that, despite selection on different hosts, *E. coli* B remained the most highly permissive host and thus was used for quantification of phage titer throughout this study.

**Figure 3:**
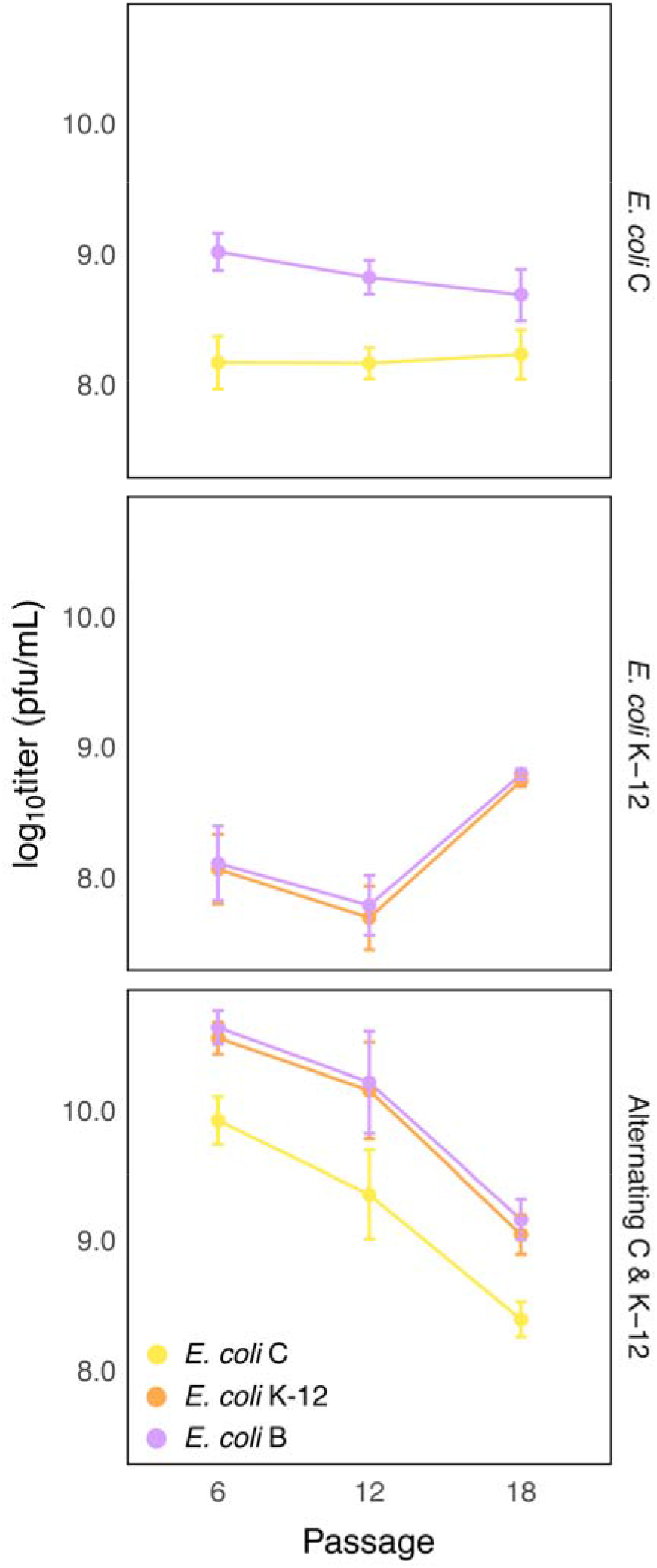
Quantification of the evolved phage populations. The 15 populations were quantified for phage titer on the original host *E. coli* B and the selection host(s) *E. coli* C and/or K-12 at passages 6, 12, 18. Plaque assays for each population was performed with threefold replication. Panels indicate one of three evolutionary histories and each point represents grand mean titer (log_10_ PFU/mL) of the five populations within each evolutionary history; error bars indicate 95% confidence limits of the grand mean. A series of *t*-tests comparing titers pooled by evolutionary history indicated that all comparisons involving *E. coli* C and B yielded significant differences (with the higher titer estimate occurring on *E. coli* B) and all comparisons involving *E. coli* K-12 and B yielded no significant differences (see main text and Table 1).

**Table 1.**
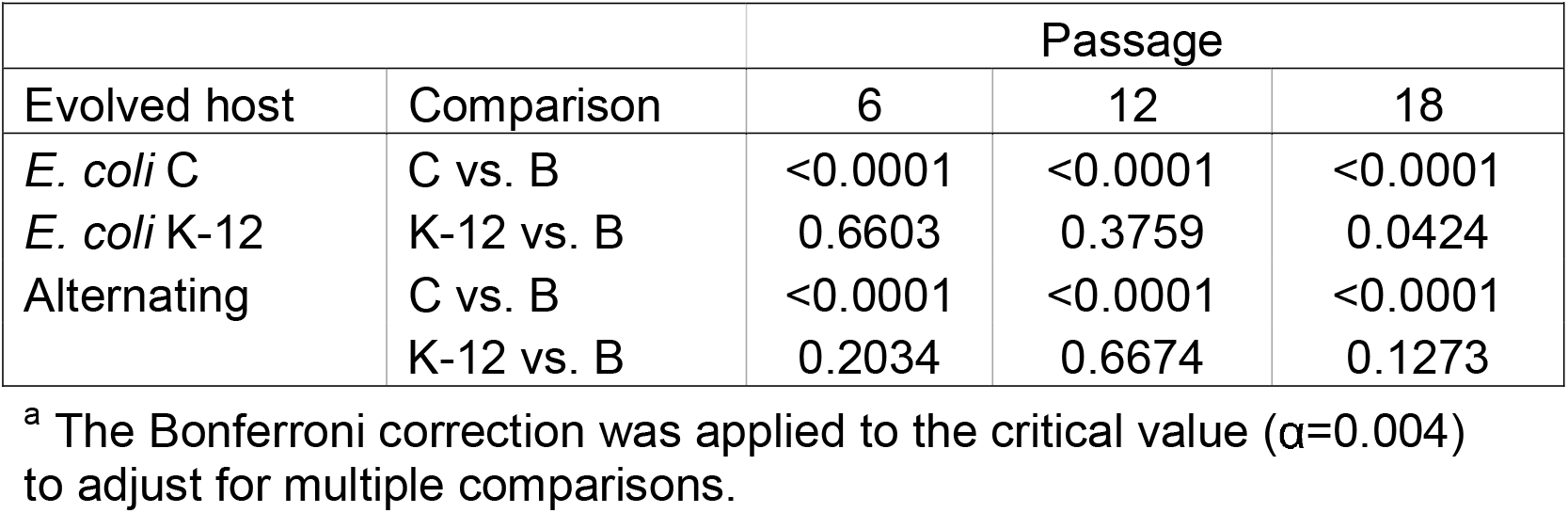
p-values^a^ of t-tests comparing log_10_ titer of T4 populations after plaque quantification on the evolved host(s) versus *E. coli* B.

### Niche-breadth evolution resulted in changes in phage productivity over time

After 20 days of host selection, the 15 evolved populations (5 x 3 replicates) were assayed for productivity on three hosts (*E. coli* C, K-12, and B) at three equidistant time points (passage 6, 12, and 18). We measured phage productivity at three time points and across histories and assay hosts, relative to the productivity of the ancestor. This experiment yielded a total of 405 productivity measurements (including *n* = 3 replicate assays). Mixed effects linear modeling fit by restricted maximum likelihood was used to test for the effects of experimental treatments—evolutionary history, assay host, and number of passages—on phage productivity. Evolutionary history, assay host, number of passages, and their interactions were specified as fixed factors and population nested within history was specified as a random factor. We used the model to generate predicted least squared (LS) means for productivity at each level of evolutionary history, assay host, and number of passages (Fig. 4). Due to the presence of interactions, a type III analysis of deviance table was calculated to determine the significance of each fixed parameter and their interactions in explaining variation in phage productivity (Table 2). This analysis revealed that all fixed factors and their interactions had a significant effect on productivity.

**Figure 4:**
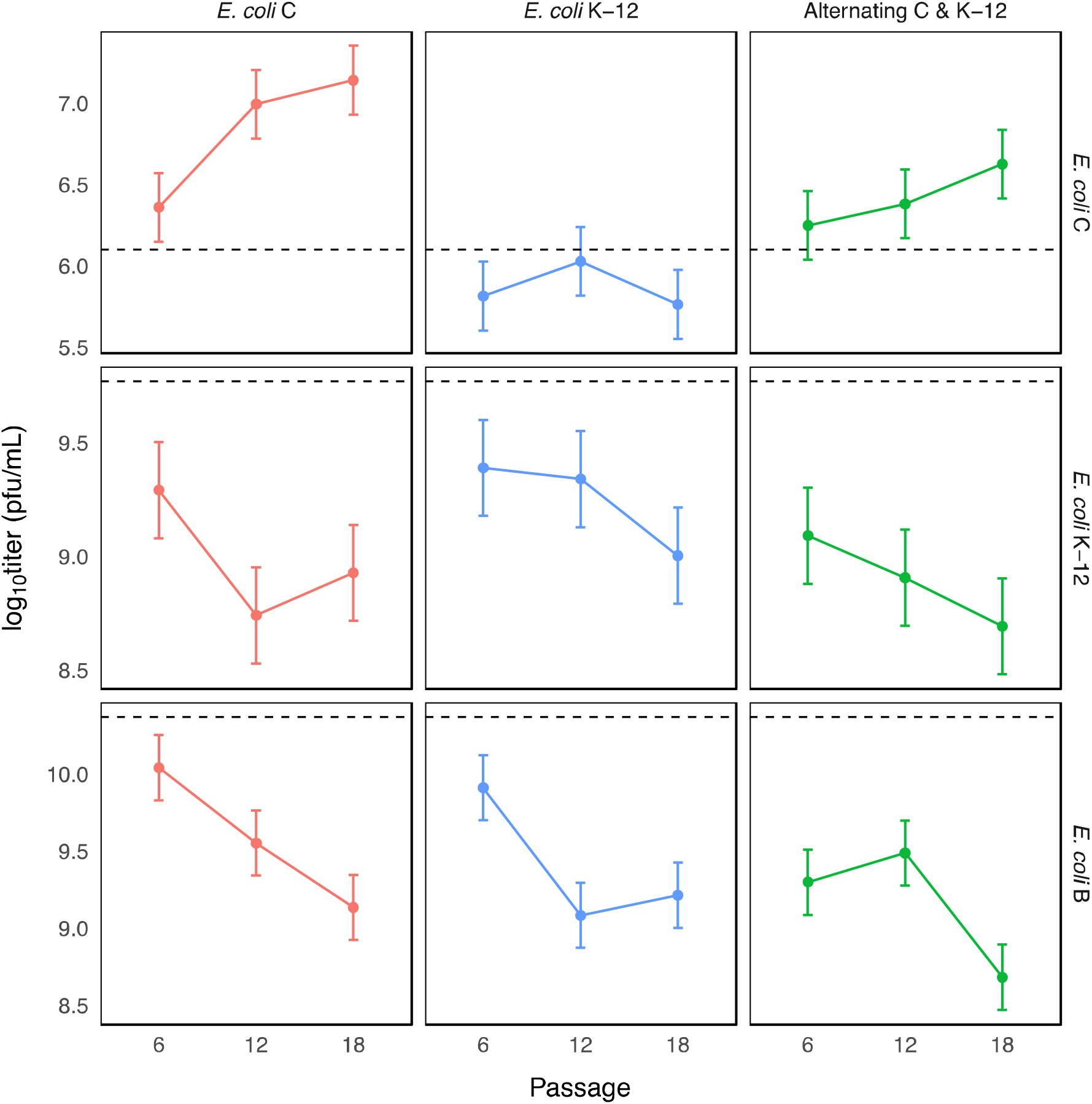
Model predictions of mean productivity. Predicted LS means for productivity at each level of evolutionary history, assay host, and number of passages (estimates were not provided for individual populations because population was specified as a random factor). Horizontal rows denote the assay host and vertical columns and colors denote evolutionary history. Each point represents LS mean productivity (log_10_ titer [pfu/mL]) and error bars indicate 95% confidence limits. Dashed black lines indicate the productivity of the T4 ancestor on a particular assay host to provide a frame of reference.

**Table 2.** Analysis of deviance table^a^ for a mixed effects linear model testing the effect of each fixed parameter and their interactions on productivity (log_10_ titer) of evolved T4 populations.

| Fixed effects | Chi-sq | Df | Pr(>Chi-sq) |
| --- | --- | --- | --- |
| Evolutionary history | 14.898 | 2 | 0.0005 |
| Assay host | 1004.209 | 2 | <0.0001 |
| Number of passages | 45.812 | 2 | <0.0001 |
| History x assay | 45.366 | 4 | <0.0001 |
| History x passages | 28.224 | 4 | <0.0001 |
| Assay x passages | 115.056 | 4 | <0.0001 |
| History x assay x passages | 74.234 | 8 | <0.0001 |

<sup>a</sup> Type III Wald chi-sq test.

In addition, we performed pairwise comparisons on predicted LS means for productivity between evolutionary histories for each level of assay host and number of passages (see Fig. 4 for a graphical representation and Supplementary Table 1a and 1b for numerical estimates). Focusing on the final time point (passage 18) and assay host *E. coli* C, all contrasts of predicted means for phage productivity between evolutionary histories were significant (*P* ≤ 0.001), indicating that by passage 18, evolutionary history led to different outcomes of productivity on *E. coli* C. Predicted productivity was highest for populations evolved on *E. coli* C (*LS mean* = 7.14, *95% CI* = [6.93; 7.35]), lower for populations evolved on alternating hosts (*LS* mean = 6.63, *95% CI* = [6.41; 6.84]), and lowest for populations evolved on *E. coli* K-12 (*LS mean* = 5.76, *95% CI* = [5.55; 5.97]). At passage 18 and assay host *E. coli* K-12, all contrasts of predicted means for phage productivity between evolutionary histories were not significant (*P* > 0.1), indicating that evolutionary history had the same effect on productivity on *E. coli* K-12 at this time point. Furthermore, all predicted productivities on *E. coli* K-12 were lower than that of the ancestor (*E. coli* C *LS mean* = 8.93, *95% CI* = [8.72; 9.14]; *E. coli* K-12 *LS mean* = 9.00, *95% CI* = [8.79; 9.21]; alternating host *LS mean* = 8.69, *95% CI* = [8.48; 8.90]). At passage 18 and assay host *E. coli* B, contrasts of predicted means for phage productivity between the alternating host history and the *E. coli* C and *E. coli* K-12 histories were significant (*P* < 0.01) while the difference between the predicted means of the two single host histories was not significant (*P* = 0.60). Interestingly, all predicted productivities on *E. coli* B were also lower than that of the ancestor, with the lowest productivity belonging to the alternating history (*E. coli* C *LS mean* = 9.14, *95% CI* = [8.92; 9.35]; *E. coli* K-12 *LS mean* = 9.21, *95% CI* = [9.00; 9.43]; alternating host *LS mean* = 8.68, *95% CI* = [8.47; 8.89]).

### Genetic signatures are suggestive of positive selection in evolved populations

Following the experimental evolution, the genomes of the 15 evolved populations at passage 18 were sequenced to analyze mutations and their frequencies. The small genome size of T4 coupled with current sequencing technology yielded over 1000× coverage depth for most positions in the genome, with over 95% of the reads mapping unambiguously to the T4 reference genome; this enabled the reliable detection of polymorphisms occurring at 5% frequency or greater. A total of 174 mutations occurred across 145 loci, with a range of 3–30 mutations per population and 77.6% occurring as unique to a particular population (for a graphical representation see Fig. 5, and for a complete list see Supplementary Table 2a). Of the total number of mutations, 2.8% occurred in intergenic regions, 6.9% were indels, 82.2% were single nucleotide polymorphisms (SNPs), and 8.1% were multi-nucleotide polymorphisms (xNPs). All xNPs were nonsynonymous and among them, 14.7% were synonymous and 85.3% were nonsynonymous. Of the total number of detected SNPs, we observed a dN/dS ratio of 5.8. If observed indels and xNPs are also classified as nonsynonymous (each instance is conservatively counted as one nonsynonymous mutation despite causing more than one amino acid change in most cases), this ratio increases to 7.0 (Figure 6 and Table 3). Across evolutionary histories, we observed that few mutations reached fixation, with the preponderance of those being nonsynonymous mutations. These signatures indicate that positive selection shaped phenotypic changes during experimental evolution.

**Table 3.**
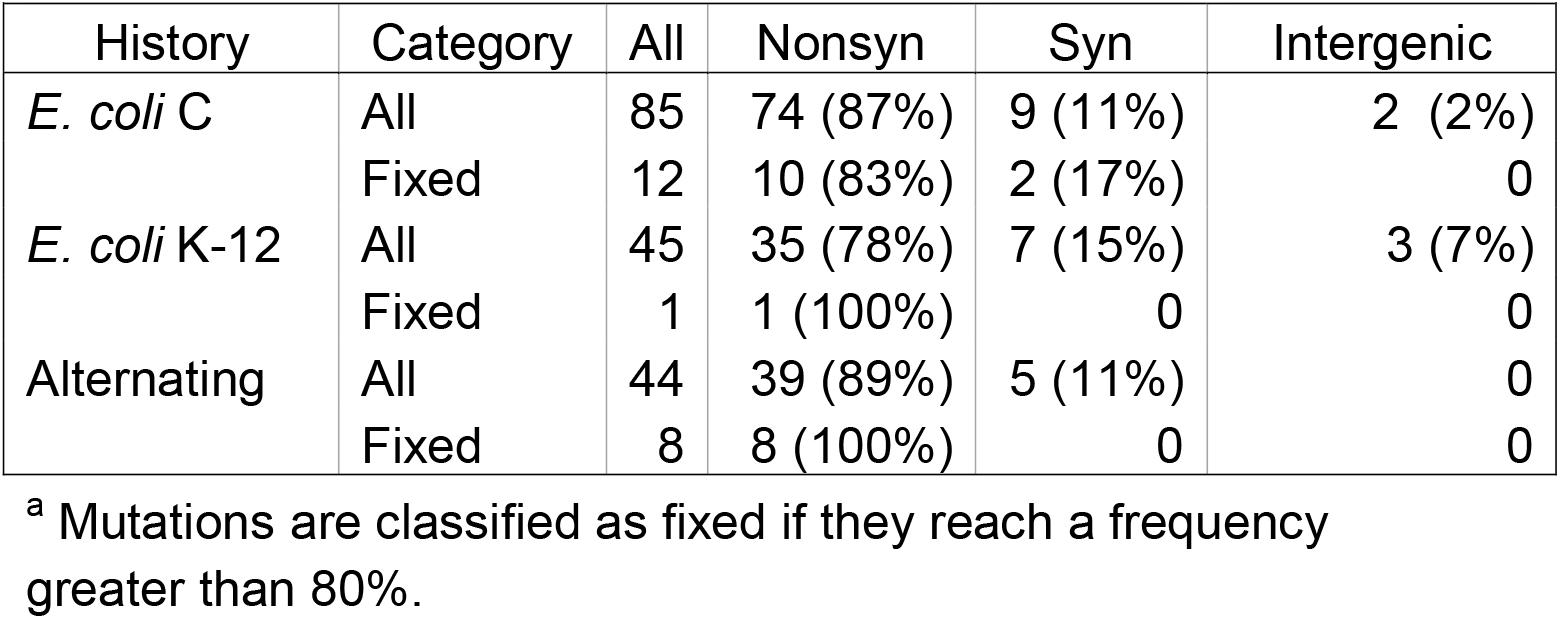
The total number of observed and fixed^a^ mutations in the 15 sequenced populations, grouped by evolutionary history and sorted by type. The percentage of mutations observed and fixed in a given type are shown in parentheses.

| History | Category | All | Nonsyn | Syn | Intergenic |
| --- | --- | --- | --- | --- | --- |
| *E. coli* C | All | 85 | 74 (87%) | 9 (11%) | 2 (2%) |
|  | Fixed | 12 | 10 (83%) | 2 (17%) | 0 |
| *E. coli* K-12 | All | 45 | 35 (78%) | 7 (15%) | 3 (7%) |
|  | Fixed | 1 | 1 (100%) | 0 | 0 |
| Alternating | All | 44 | 39 (89%) | 5 (11%) | 0 |
|  | Fixed | 8 | 8 (100%) | 0 | 0 |

<sup>a</sup> Mutations are classified as fixed if they reach a frequency greater than 80%.

**Figure 5.**
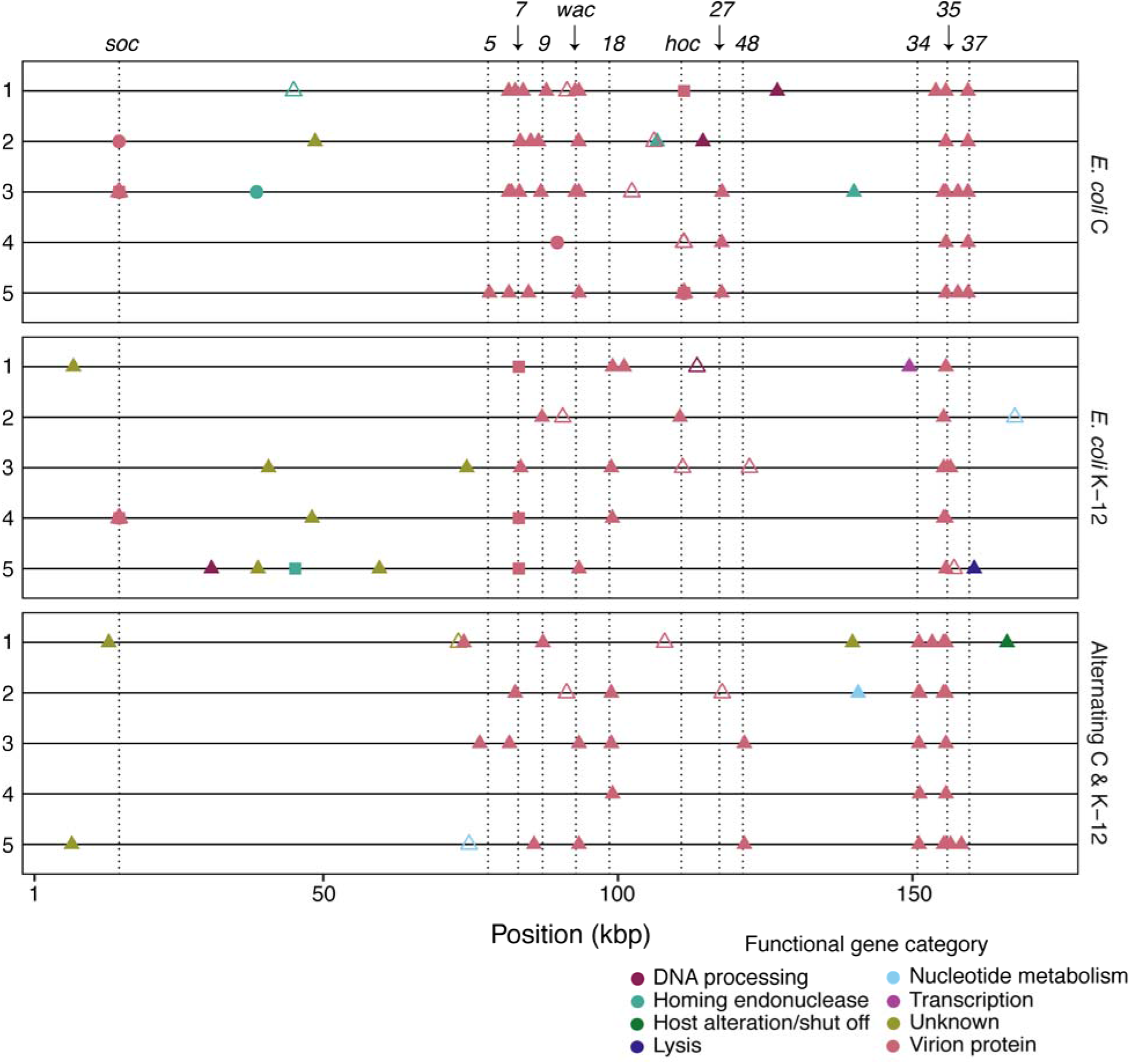
The distribution of new mutations detected in the evolved phage populations. Panels indicate one of three evolutionary histories and lines within panels represent particular populations within an evolutionary history; numbers for each line correspond with the numbered populations from Figure 4. Each point represents a new mutation (relative to the ancestor) detected at ≥ 5% frequency. Point shape indicates type of mutation (filled triangles: non-synonymous; empty triangles: synonymous; circles: indel; squares: xNPs; diamonds: intergenic). Colors indicate functional gene category; labels above the top panel and dashed lines denote relative positions of virion structural genes.

**Figure 6.**
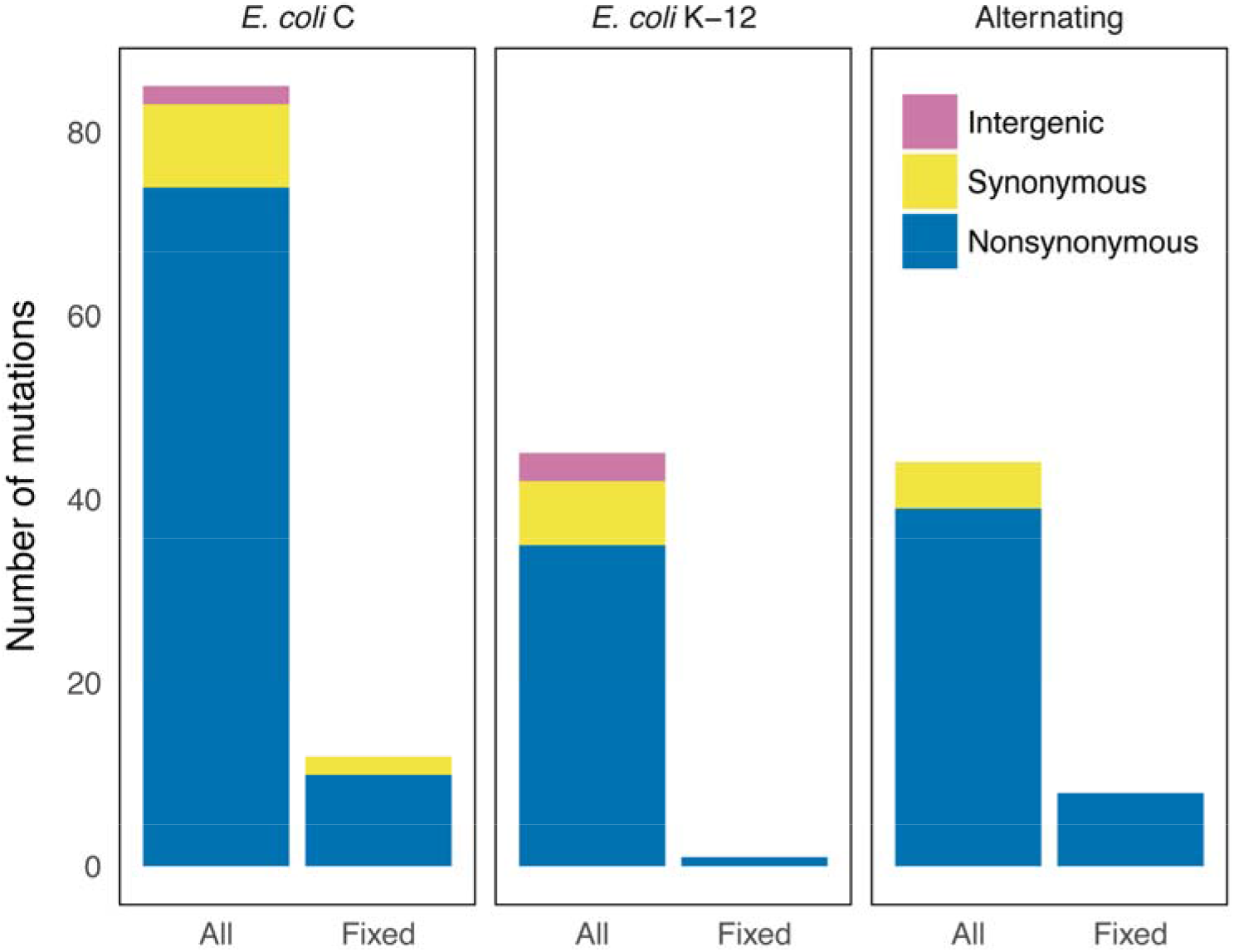
Classification of mutations. Panels indicate one of three histories; categories within panels indicate the total number of observed (>5%) and fixed (>80%) mutations for the five parallel populations within each history. Bar colors represent the type of mutation for each category: intergenic, synonymous, and nonsynonymous.

### Mutations in structural genes were predominant in evolved T4 populations

Mutations during evolution tended to cluster in regions of the T4 genome containing genes that code for structural proteins. Furthermore, of the 62 polymorphisms that reached a frequency of 50% or greater, all but four occurred in structural genes. Following these observations, a Scheirer-Ray-Hare (SRH) non-parametric two-way ANOVA (Sokal & Rohlf, 2011) was performed to formally test whether different regions of the genome experienced different rates of mutation, and if the rates also depended on the evolutionary history of the population. First, the number of nucleotide sites that each functional category occupies in the genome was then determined. From this, the mutation rate (number of mutations per nucleotide site) of each category was calculated for the 15 evolved populations. Evolutionary history, functional category, and their interaction were specified as factors in the SRH non-parametric two-way ANOVA. This analysis showed that the rate of mutation was different across functional gene categories (*P* < 0.001). There was neither a significant effect for evolutionary history (*P* = 0.588) nor for interactions between functional category and history (*P* = 0.789). To follow up, pairwise comparisons were generated using Dunn’s test, which revealed that only comparisons involving the structural gene category were significant (*P* < 0.01). Specifically, this indicated that the mutation rate of the structural gene category was significantly higher (i.e., mutations were overrepresented in structural genes) than all other categories.

## Discussion

In this study, we performed experimental evolution using *Escherichia virus T4* to examine the phenotypic and molecular manifestations of niche-breadth evolution. Our findings demonstrate that niche-breadth evolution led to measurable changes in phage productivity. For populations grown on *E. coli* C, productivity on *E. coli* C increased while productivity decreased on both *E. coli* B and K-12, indicating that adaptation on *E. coli* C led to reduced performance on both the original and unselected hosts. These results conform to theoretical expectations where evolved populations improve performance in the selective environment at the expense of diminished performance in alternate environments. For the alternating host evolved populations, the magnitude of increased productivity on *E. coli* C was smaller than that of the populations that only experienced the same host, which suggests that exposure to two hosts and temporal variability limited adaptation on *E. coli* C. This result also conforms to theoretical expectations; specialists are predicted to evolve faster than generalists because the time to fixation of favorable alleles is shorter for specialists (Whitlock, 1996).

### Experimental populations deviated from the expectations of simple tradeoff models

The observed changes in productivity on *E. coli* K-12 for the *E. coli* K-12 and alternating host populations offered surprising results. For example, these populations decreased in productivity on *E. coli* K-12, despite being experimentally exposed to this host. These results were not only unforeseen, they ostensibly deviate from theoretical expectations. We predicted that evolution on *E. coli* K-12 would lead to higher productivity with the single host populations performing better than the alternating host populations. Instead, we observed decreased productivity with no significant differences between the single and alternating host histories. Though empirical evidence from previous viral experimental evolution studies have contradicted theory and found no cost associated with generalism (Bedhomme et al., 2012; Novella et al., 1999; Turner & Elena, 2000), our results present a different kind of departure. This raises an important question specific to our system: why would T4 passaged on *E. coli* K-12 evolve lower productivity on its selective host? One simple explanation is that productivity did not improve because it was not the target of selection, which undermines the neat equivalence between productivity and reproductive fitness. In contexts where competition is more synonymous with fitness, one might expect productivity to be completely decoupled from evolution. Indeed, prior studies have suggested that productivity can be negatively correlated with competitive ability in T4 in the setting of experimental evolution (Kerr et al., 2006).

### Study limitations

As with any exercise in experimental evolution used to investigate an ecological phenomenon, there are many limitations that can affect our interpretations and conclusions. The chief concern is that measurements made in the laboratory may not be relevant to organisms in their natural context. However contrived, evolutionary and ecological phenomena still apply to our experimental microcosm, which makes it nevertheless useful for examining ecological questions. Consequently, we are comfortable interpreting our results and discussing them in light of broader ecological theory. Next, our study did not measure changes in phage fitness, but rather, phage productivity. Though this trait can be closely associated with phage fitness and has been previously used as a proxy for fitness (Morley et al., 2015; Turner, Draghi, & Wilpiszeski, 2012), it is clear from our findings that assuming this equivalence is not always appropriate in our system. Importantly, this suggests that the productivity to fitness equivalence might not exist for other virus-host systems. Lastly, our interpretations should consider the potentially complicating role of phenotypic plasticity in these results, as phage phenotypes may be moderated by the host encountered. For example, previous work in T4 has shown that expression of T4 mutant phenotypes can be moderated by bacterial host strain (Benzer, 1957).

### Evolutionary genomics data reveals evidence for selection

We observed genetic signatures that are suggestive of selection shaping the evolved populations during niche breadth evolution. Specifically, the high dN/dS ratio, along with the observation that few nonsynonymous mutations reached fixation, indicates selection.

The genotypic evidence is suggestive of positive selection as the source of elevated rate of nonsynonymous mutations in structural genes (for a more detailed discussion of other plausible causes, such as mutation bias, see the Supplemental Information). Though mutation bias can influence the direction of an adaptive trajectory, newly introduced mutations are still subject to elimination or fixation through the processes of drift and selection. As such, it is probable that fixed (or high frequency) nonsynonymous mutations are of adaptive significance.

### The role of structural genes in bacteriophage niche-breadth evolution

Our findings demonstrate that structural genes are important for niche-breadth evolution: i) the overwhelming majority (82%) of new mutations occurred in structural genes, ii) nearly all (94%) high frequency mutations (>50%) occurred in structural genes, and iii) we detected a significantly higher mutation rate in structural genes. Notably, these rates did not differ across evolutionary histories, which indicates that structural genes are important niche-breadth evolution, regardless of the host(s) encountered and level of temporal variability of the environment. Structural genes encode for four different types of virion proteins: capsid, neck, tail, and tail fibers (Miller et al., 2003). Of the 142 new mutations detected in structural genes, 44% occurred in genes coding for tail fiber proteins. We observed mutations in gene *37*, which encodes the distal subunit of the LTF (gp37) that directly interacts with bacterial receptors (Bartual et al., 2010). This supports previous evidence suggesting that mutation in gene *37* can alter the niche breadth of T4 (Tétart, Repoila, Monod, & Krisch, 1996). Interestingly, the majority of mutations (74%) detected in LTF genes occurred in the genes encoding the LTF proximal subunits (i.e., gp34 and gp35). This suggests that structures that are only indirectly responsible for host recognition are also important to niche-breadth evolution. Mutations in tail genes accounted for 30% of structural mutations; the great majority of those mutations (82%) occurred in genes that code for various baseplate components (e.g., genes *5-10*, *27*, *48*, and *54*). Mutations in baseplate genes occurred across all three evolutionary histories, in nearly every population (14 out of 15). This suggests that the baseplate, which regulates infection (Yap et al., 2016), may serve an important role in the evolution of niche breath. The remaining structural mutations were detected in six genes that code for capsid proteins, with the vast majority (90%) occurring in two of those six genes: *soc* (small outer capsid) and *hoc* (head outer capsid). This suggests that soc and hoc proteins, which provide stability to the capsid (Fokine et al., 2004), may also be important to niche breadth evolution. For those interested in a more detailed discussion of mutations, e.g. parallel substitutions or distribution across populations, please refer to the Supplemental Discussion.

## Conclusions

In this study, we used experimental evolution of *Escherichia virus T4* to characterize the phenotypic and molecular changes associated with niche breadth evolution. Our findings indicate that populations evolved measurable and meaningful changes in phage productivity. These phenotypic results confirmed some and contradicted other theoretical expectations, which demonstrates the complexity of niche-breadth evolution.

Genomic sequencing of evolved populations enabled us to detect a significantly higher rate of mutation in structural genes. Notably, mutation rates of structural genes were similar across evolutionary histories, which indicates that selection in structural genes is important to niche-breadth evolution, regardless of conditions that promoted a particular ecological strategy (specialism or generalism). Our study presents compelling evidence that structural genes serve an important role in how phage evolve different niche breadth strategies, a finding with many implications and broad applications.

## Acknowledgements

JYP was supported by the National Science Foundation Graduate Research Fellowship Program under Grant No. DGE1745303. The authors would like to thank Kalsang Namgyal for help with media preparation and serial propagation, Robin Hopkins, Elizabeth Wolkovich, and Steven Worthington for consultation on statistical approaches. The authors would like to thank Tim Sackton for advice on sequencing and linear models, Claire Reardon for advice on library preparation and sequencing, and Michael Desai for providing laboratory space and reagents for library preparation. CBO acknowledges funding support from NSF RII Track-2 FEC award number 1736253.

## Data Accessibility

Sequence data will be deposited to the NCBI Sequence Read Archive. Phenotypic data will be archived and made publicly accessible via Dryad.

## Author Contributions

JYP designed the research, executed experiments, conducted statistical analyses, and authored the manuscript. CBO and DLH designed the research, provided technical guidance, and coauthored the manuscript. ANNB provided guidance on library preparation, performed read analysis, and coauthored the manuscript.

## References

Abedon, S. T., Herschler, T. D., & Stopar, D. (2001). Bacteriophage Latent-Period Evolution as a Response to Resource Availability. Applied and Environmental Microbiology, 67(9), 4233–4241. doi:10.1128/aem.67.9.4233-4241.2001

Bartual, S. G., Otero, J. M., Garcia-Doval, C., Llamas-Saiz, A. L., Kahn, R., Fox, G. C., & van Raaij, M. J. (2010). Structure of the bacteriophage T4 long tail fiber receptor-binding tip. Proc Natl Acad Sci U S A, 107(47), 20287–20292. doi:10.1073/pnas.1011218107

Bates, D., Mächler, M., Bolker, B., & Walker, S. (2015). Fitting Linear Mixed-Effects Models Using lme4. Journal of Statistical Software, 67(1). doi:10.18637/jss.v067.i01

Baym, M., Kryazhimskiy, S., Lieberman, T. D., Chung, H., Desai, M. M., & Kishony, R. (2015). Inexpensive multiplexed library preparation for megabase-sized genomes. PLoS One, 10(5), 1–15.

Bedhomme, S., Lafforgue, G., & Elena, S. F. (2012). Multihost experimental evolution of a plant RNA virus reveals local adaptation and host-specific mutations. Mol Biol Evol, 29(5), 1481–1492. doi:10.1093/molbev/msr314

Benzer, S. (1957). The elementary units of heredity. The elementary units of heredity.

Bolger, A. M., Lohse, M., & Usadel, B. (2014). Trimmomatic: a flexible trimmer for Illumina sequence data. Bioinformatics, 30(15), 2114–2120. doi:10.1093/bioinformatics/btu170

Cooper, L. A., & Scott, T. W. (2001). Differential evolution of eastern equine encephalitis virus populations in response to host cell type. Genetics, 157(4), 1403–1412.

Crill, W. D., Wichman, H. A., & Bull, J. (2000). Evolutionary reversals during viral adaptation to alternating hosts. Genetics, 154(1), 27–37.

Deatherage, D. E., & Barrick, J. E. (2014). Identification of mutations in laboratory-evolved microbes from next-generation sequencing data using breseq. Methods Mol Biol, 1151, 165–188. doi:10.1007/978-1-4939-0554-6_12

Demerec, M., & Fano, U. (1944). Bacteriophage-resistant mutants in Escherichia coli. Genetics, 19, 119–136.

Duffy, S., Burch, C. L., & Turner, P. E. (2007). Evolution of host specificity drives reproductive isolation among RNA viruses. Evolution, 61(11), 2614–2622. doi:10.1111/j.1558-5646.2007.00226.x

Duffy, S., Turner, P. E., & Burch, C. L. (2006). Pleiotropic costs of niche expansion in the RNA bacteriophage ϕ6. Genetics, 172(2), 751–757.

Fokine, A., Chipman, P. R., Leiman, P. G., Mesyanzhinov, V. V., Rao, V. B., & Rossmann, M. G. (2004). Molecular architecture of the prolate head of bacteriophage T4. Proceedings of the National Academy of Sciences of the United States of America, 101(16), 6003–6008.

Fox, J., & Weisberg, S. (2011). An R Companion to Applied Regression (Second ed.). Thousand Oaks, CA: Sage.

Futuyma, D. J., & Moreno, G. (1988). The evolution of ecological specialization. Annual Review of Ecology and Systematics, 19(1), 207–233.

Hadas, H., Einav, M., Fishov, I., & Zaritsky, A. (1997). Bacteriphage T4 development depends on the physiology of its host Escherichia coli. Microbiology, 143, 179–185.

Kassen, R. (2002). The experimental evolution of specialists, generalists, and the maintenance of diversity. Journal of evolutionary biology, 15(2), 173–190.

Kawecki, T. J. (1994). Accumulation of Deleterious Mutations and the Evolutionary Cost of Being a Generalist. The American Naturalist, 144(5), 833–838.

Kerr, B., Neuhauser, C., Bohannan, B. J., & Dean, A. M. (2006). Local migration promotes competitive restraint in a host-pathogen ‘tragedy of the commons’. Nature, 442(7098), 75–78. doi:10.1038/nature04864

Kutnjak, D., Elena, S. F., & Ravnikar, M. (2017). Time-Sampled Population Sequencing Reveals the Interplay of Selection and Genetic Drift in Experimental Evolution of Potato Virus Y. J Virol, 91(16). doi:10.1128/JVI.00690-17

Lenski, R. E., Rose, M. R., Simpson, S. C., & Tadler, S. C. (1991). Long-Term Experimental Evolution in Escherichia coli. I. Adaptation and Divergence Durng 2,000 Generations. American Society of Naturalists., 138(6), 1315–1341.

Lenth, R. V. (2016). Least-Squares Means: The R Package lsmeans. Journal of Statistical Software, 69(1). doi:10.18637/jss.v069.i01

Levins, R. (1968). Evolution in changing environments: some theoretical explorations: Princeton University Press.

Lynch, M., & Gabriel, W. (1987). Environmental tolerance. The American Naturalist, 129(2), 283–303.

MacLean, R. C., Bell, G., & Rainey, P. B. (2004). The evolution of a pleiotropic fitness tradeoff in Pseudomonas fluorescens. Proceedings of the National Academy of Sciences of the United States of America, 101(21), 8072–8077.

Mangiafico, S. (2018). rcompanion: Functions to Support Extension Education Program Evaluation. R package version 1.11.3.

Miller, E. S., Kutter, E., Mosig, G., Arisaka, F., Kunisawa, T., & Ruger, W. (2003). Bacteriophage T4 Genome. Microbiology and Molecular Biology Reviews, 67(1), 86–156. doi:10.1128/mmbr.67.1.86-156.2003

Miralles, R., Moya, A., & Elena, S. F. (2000). Diminishing Returns of Population Size in the Rate of RNA Virus Adaptation. Journal of Virology, 74(8), 3566–3571.

Monod, J. (1949). The growth of bacterial cultures. Annual Reviews in Microbiology, 3(1), 371–394.

Morley, V. J., Mendiola, S. Y., & Turner, P. E. (2015). Rate of novel host invasion affects adaptability of evolving RNA virus lineages. Proc. R. Soc. B, 282(1813), 20150801.

Novella, I. S., Clarke, D. K., Quer, J., Duarte, E. A., Lee, C. H., Weaver, S. C., … Holland, J. J. (1995). Extreme fitness differences in mammalian and insect hosts after continuous replication of vesicular stomatitis virus in sandfly cells. Journal of Virology, 69(11), 6805–6809.

Novella, I. S., Hershey, C. L., Escarmis, C., Domingo, E., & Holland, J. J. (1999). Lack of evolutionary stasis during alternating replication of an arbovirus in insect and mammalian cells. Journal of molecular biology, 287(3), 459–465.

Ogle, D. H. (2018). FSA: Fisheries Stock Analysis. R package version 0.8.19.

Ogbunugafor, C.B, et al. “Evolution of increased survival in RNA viruses specialized on cancer-derived cells.” The American Naturalist 181.5 (2013): 585–595

Palaima, A. (2007). The fitness cost of generalization: present limitations and future possible solutions. Biological journal of the linnean society, 90(4), 583–590.

R Development Core Team. (2017). R: A language and environment for statistical computing. Vienna, Austria: R Foundation for Statistical Computing.

Sexton, J. P., Montiel, J., Shay, J. E., Stephens, M. R., & Slatyer, R. A. (2017). Evolution of ecological niche breadth. Annual Review of Ecology, Evolution, and Systematics, 48.

Sokal, R. R., & Rohlf, F. J. (2011). Biometry: the principles and practice of statistics in biological research (4th ed.). New York: W. H. Freeman and Co.

Tétart, F., Repoila, F., Monod, C., & Krisch, H. (1996). Bacteriophage T4 host range is expanded by duplications of a small domain of the tail fiber adhesin. In: Elsevier.

Turner, P. E., Draghi, J. A., & Wilpiszeski, R. (2012). High-throughput analysis of growth differences among phage strains. J Microbiol Methods, 88(1), 117–121. doi:10.1016/j.mimet.2011.10.020

Turner, P. E., & Elena, S. F. (2000). Cost of Host Radiation in an RNA Virus. Genetics, 156, 1465–1470.

Whitlock, M. C. (1996). The Red Queen Beats the Jack-of-all-Trades: The Limitations on the Evolution of Phenotypic Plasticity and Niche Breadth. The American Naturalist, 148, S65–S77.

Wichman, H. A., Millstein, J., & Bull, J. J. (2005). Adaptive molecular evolution for 13,000 phage generations: a possible arms race. Genetics, 170(1), 19–31. doi:10.1534/genetics.104.034488

Wilson, D. S., & Yoshimura, J. (1994). On the Coexistence of Specialists and Generalists. The American Naturalist, 144(4), 692–707.

Yap, M. L., Klose, T., Arisaka, F., Speir, J. A., Veesler, D., Fokine, A., & Rossmann, M. G. (2016). Role of bacteriophage T4 baseplate in regulating assembly and infection. Proc Natl Acad Sci U S A, 113(10), 2654–2659. doi:10.1073/pnas.1601654113

Yu, F., & Mizushima, S. (1982). Roles of lipopolysaccharide and outer membrane protein OmpC of Escherichia coli K-12 in the receptor function for bacteriophage T4. Journal of bacteriology, 151(2), 718–722.

Zuur, A. F., Ieno, E. N., & Elphick, C. S. (2010). A protocol for data exploration to avoid common statistical problems. Methods in Ecology and Evolution, 1(1), 3–14. doi:10.1111/j.2041-210X.2009.00001.x

