## Supplemental Information for "Experimental evolution for niche breadth in bacteriophage T4 highlights the importance of structural genes"

### Bacteriophage T4 structural genes evolve rapidly during experimental evolution for niche breadth expansion

Here we offer extended discussion points and analyses related to the findings presented in the main text. In particular, this section provides additional detail on several of the molecular findings presented in the main text.

**Notes on analysis of mutations located in capsid genes.** The majority of mutations in outer capsid genes were detected in the *hoc* gene. In particular, one *E. coli* C evolved lineage sustained 24 *hoc* mutations that occurred at nearly equivalent frequencies (53-57%), and were observed in nucleotide positions which were in close proximity or adjacent to each other. Visualization of sequencing reads containing the relevant positions revealed that these polymorphisms appeared as a block (reads contained all or none of the mutations), which indicates a major but singular mutational event. A second set of polymorphisms detected at approximately 77% frequency was observed in a different *E. coli* C evolved lineage. This six base-pair mutation resulted in two amino acid substitutions; again, visualization of reads revealed that these mutations appeared as a block. These observations are particularly interesting in light of functional characterizations of *soc* and *hoc*. Viable T4 mutants missing either or both proteins have been isolated, demonstrating that these proteins are nonessential for phage growth (Ishii & Yanagida, 1977). In fact, the dispensable nature of these proteins provided the basis for T4-phage display, a technique in which novel protein and polypeptide sequences are inserted into *soc* and/or *hoc*, resulting in display on the phage surface for identification and characterization (Ren & Black, 1998). However, soc proteins are known to provide stability to the capsid in extreme pH and temperatures, whereas hoc proteins have a more marginal effect on stability (Andrei Fokine et al., 2004). This indicates that the mutations we observed are unlikely to be severely deleterious, but it is presently unclear if they are selectively neutral or beneficial. However, the two *E. coli* C evolved lineages which sustained these mutations were also the lineages that experienced the greatest magnitude of productivity increase on *E. coli* C (see lineages 1 and 5 of Fig. 4). Furthermore, lineage 5 of the *E. coli* C history sustained two additional fixed nonsynonymous mutations in the *hoc* gene. These observations are possibly a coincidence. Conversely, they could indicate a potential beneficial effect (e.g., increased virion stability or rate of capsid morphogenesis).

The adsorption of T4 on *E. coli* is initiated by the tail fibers, beginning with a reversible interaction between the long tail fibers (LTF) and receptors on the bacterial cell surface (lipopolysaccharide and/or OmpC proteins), followed by irreversible binding of the short tail fibers (STF) (Kostyuchenko et al., 1999; Yu & Mizushima, 1982). The T4 tail fiber proteins are thus responsible for interactions that are the gatekeepers of infection;

following adhesion, phage DNA is injected into the bacterium for replication and subsequent completion of the lytic life cycle. The LTF is a multiprotein complex composed of four different gene products: gp34, gp35, gp36, and gp37. Gp36 and gp37 are the subunits that form the distal half of the LTF, with gp37 acting as the portion of the LTF that is directly involved with bacterial receptor recognition. Gp34 and gp35 are the subunits that form the proximal half of the LTF, with gp35 forming the “kneecap” hinge and gp34 interacting with the phage baseplate (Bartual et al., 2010; Leiman et al., 2010).

Previous work has shown that duplications in gene *37* (encodes gp37) expands the T4 host range to include *Yersinia*, a bacterial genus that wild-type laboratory strains of T4 cannot infect (Tétart, Repoila, Monod, & Krisch, 1996). We observed mutations in genes *12* (encodes STF) and *37*; however, all three mutations observed in *12* were synonymous, and only present in three lineages at frequencies < 13%, while eight of the nine mutations observed in *37* were non-synonymous and occurred in seven lineages at frequencies between 5–100%, across all three histories. Notably, one codon position in *37* was substituted with two different amino acid residues: Y953H and Y953F; these substitutions only occurred in the *E. coli* C history, with the first detected in three lineages and the latter detected in two lineages (Supplementary Table 2a). The Y953H mutation was fixed across the three lineages in which it was detected while the Y953F mutation occurred at frequencies of 17% and 25%. It is possible that these mutations may be specifically adaptive to *E. coli* C because replacements at this particular position only occurred in one evolutionary history.

One particular nonsynonymous mutation (G323D) in gene *35* was detected in 12 of the 15 lineages. Two of the three lineages without this mutation sustained a different nonsynonymous substitution (T193M) in gene *35* and three lineages displayed both substitutions. The G323D substitution was fixed in four lineages (two in the alternating history and one in each single host history) and detected at high frequency (≥ 60%) in five other lineages (across three histories). The high frequencies and degree of parallelism of the G323D substitution is striking and suggests an interplay between mutation bias and selection. Given the interaction between LTF proteins, one might hypothesize that mutations in gene *35* are compensatory to mutations in gene *37*, but the presence of eight lineages with polymorphisms detected in gene *35* and not gene *37* challenges that hypothesis. Rather, given the distribution and high frequency of G323D observed across all three histories, we hypothesize that this substitution is universally beneficial to phage growth capabilities.

In the case of gene *34*, a different pattern emerged: there were no parallel mutations, however, nearly all of the mutations in this gene (8 out of 9) were sustained by lineages evolved in the alternating host environment. Different substitutions at the same codon position were detected in four different lineages. In two of those lineages, the substitutions A89T and A89V were respectively observed at frequencies of 7% and

51%. Separately, the substitutions G133S and G133R were observed in two other lineages; the first amino acid replacement reached a frequency of 76% while the latter became fixed in the population. These observations indicate the possibility that mutations of gene *34* are of adaptive significance to a temporally variable host environment (i.e., generalist strategy).

Another nonsynonymous mutation (R417H) that displayed high frequencies with parallelism (6 out of 15 lineages, across three histories) occurred in the *wac* gene. The R417H substitution reached fixation in one lineage and high frequency (≥ 60%) in a second lineage (frequencies in other lineages were between 6–41%). Furthermore, we observed other fixed substitutions in *wac*: V207E in one *E. coli* C evolved lineage and V414D in one alternating host evolved lineage. The gene product gpwac forms six whisker fibers that radiate from the phage neck, in manner akin to a “collar”. Gpwac is multifunctional; during morphogenesis, the whiskers facilitate the binding of LTF to the phage baseplate. In extracellular conditions, gpwac performs an important regulatory function: the whiskers bind the LTF, holding them in a retracted position, thus acting as a rudimentary environmental sensor that prevents adsorption in conditions unfavorable for infection (*e.g.*, low pH) (A. Fokine et al., 2013; Letarov, Manival, Desplats, & Krisch, 2005; Terzaghi, Terzaghi, & Coombs, 1979; Wood, Eiserling, & Crowther, 1994). Mutations in *wac* may be beneficial, however, it is unclear how the traits that are governed by gpwac were affected. It is conceivable that mutations in the LTF (along with mutations in the baseplate, which were also observed) required accommodation during morphogenesis, though it is also possible that the capacity of environmental sensing was affected.

Of the mutations that occurred in tail genes, 82% occurred in genes that code for various baseplate components (*5-10*, *27*, *48*, and *54*). In total, three nonsynonymous mutations and one 12 bp indel in baseplate genes reached fixation (all occurred separately in different lineages). Additionally, mutations in baseplate genes occurred across all three evolutionary histories in nearly every lineage (14 out of 15). The phage baseplate is a complex multiprotein structure that is attached to the contractile tail. The LTF and STF are attached to the baseplate, which changes conformation along with the irreversible binding of STF (Kostyuchenko et al., 2003). This event triggers tail sheath contraction, which drives the tail tube into the bacterial inner membrane, where phage genomic DNA is injected into the host cytoplasm. Thus, the baseplate can be regarded as the nerve center of T4, transmitting signals from the tail fibers to the phage capsid for release of DNA (Yap et al., 2016). Given the importance of the baseplate in the coordination of events that lead to infection, it is possible that mutations in baseplate genes were either compensatory to mutations in the LTF or provided an unrelated selective benefit.

We emphasize that the relationships discussed here between detected polymorphisms and changes in host-use and phage growth are plausible but nevertheless speculative. The observations do provide reasonable hypotheses and justifications to further explore the effects of these mutations through site-directed mutagenesis experiments to determine whether these mutational effects are of adaptive significance to a particular host or provide universal benefit in phage growth capabilities. Although the discussion focused on particular mutations of interest, it is possible that the effects of these mutations were modulated by epistatic interactions with other polymorphisms.

**On the possibility for mutation bias in evolved populations**. The overrepresentation of new synonymous polymorphisms in structural genes can imply the presence of mutation bias and/or selection for synonymous mutations. Previous work has demonstrated that synonymous substitutions can have non-neutral effects in RNA viruses (Bedhomme et al., 2012; Carrasco, de la Iglesia, & Elena, 2007; Coleman et al., 2008). However, site-directed mutagenesis indicates that selection at synonymous sites is very weak in DNA viruses (Cuevas, Domingo-Calap, & Sanjuan, 2012). In our study, the observed transition:transversion ratio significantly exceeded the null expectation; when this observation is considered with the aforementioned evidence, it seems unlikely that selection for synonymous mutations occurred. Below we describe how mutation bias was measured and evaluate the evidence for it in the study populations.

**Commented [PJY1]:** I really like the idea of removing mutation bias from the main manuscript altogether. The main point of structural genes being a driver of niche breadth is very clear now.

**Analysis of mutation bias.** All polymorphisms ≥ 5% that resulted in base-pair substitutions were considered in this analysis. The transition:transversion ratio of paths was calculated by tabulating the number of substitutions that occurred. The transition:transversion ratio of events was calculated by tabulating the number of instances for each substitution. For example, the substitution CGC→CAC represents one path but six events because it was observed in six populations. Depending on model details, the expected ratio of transition:transversion mutations can range between 0.4–0.5 (Stoltzfus & McCandlish, 2017). In the simplest scenario, the null expectation is 0.5 because every nucleotide is subject to one transition and two transversions. We chose the estimate of 0.5 because it is the most conservative. For the purposes of statistical testing, the transition:transversion ratio was calculated separately for each evolved population. A Pearson's chi-square test was performed to determine if the observed transition:transversion ratios differed from the null expectation. This analysis was performed in Microsoft Excel version 15.20.  it is important to note that in the absence of mutation accumulation experiments, we cannot fully conclude that there is true mutation bias.

**Evidence for mutation bias.** We further explored the processes responsible for the overrepresentation of mutations in structural genes by repeating the SRH non-parametric two-way ANOVA separately on synonymous and nonsynonymous mutations. In this analysis, we tested whether different regions of the genome experienced different rates of synonymous or non-synonymous mutation, and if the rates also depended on the evolutionary history of the population. For synonymous mutations, the rate of mutation was different across functional gene categories ($P < 0.001$) with no significant effect due to evolutionary history ($P = 0.89$) nor the interaction between functional category and history ($P = 0.73$). Pairwise comparisons indicated that the rate of synonymous mutation was significantly higher in the structural gene category ($P < 0.001$). For nonsynonymous mutations, the rate of mutation was again different across functional categories ($P < 0.001$) with no significant effect due to history ($P = 0.62$) nor the interaction of history and functional category ($P = 0.94$). Pairwise comparisons indicated that the rate of nonsynonymous mutation was significantly higher in the structural gene category ($P < 0.01$). If we assume that synonymous mutations are neutral, and thus escape the effects of selection, then the signature of elevated synonymous mutation in structural genes must be accounted for by an alternative phenomenon. One possible explanation is the presence of mutation bias (i.e., intrinsic bias in the introduction of genetic variation).

The presence of transition:transversion mutational bias has been documented in experimental microbial populations, with sequence comparisons yielding estimates that are typically 2 to 4-fold greater than null expectations (Stoltzfus & McCandlish, 2017). In this study, the transition:transversion ratio was 1.2 for paths (2-fold higher than expected) and 1.4 for events (3-fold higher). A Pearson's chi-square test was performed to determine whether our observed transition:transversion ratio was different from the null expectation; the test indicated that the observed and expected transition:transversion ratios were significantly different ($\chi^2 = 53.5$; $P < 0.001$; see "Materials and Methods" for model details).

**On the possible presence of recombination in the evolved populations.** Previous work in *Saccharomyces cerevisiae* indicates that sex alters molecular signatures of adaptation (McDonald, Rice, & Desai, 2016). In asexual populations, synonymous, nonsynonymous, and intergenic mutations are equally likely to fix because selection cannot efficiently distinguish between their effects due to hitchhiking. In contrast, selection is more efficient in sexual populations, which results in fewer fixed mutations with the overwhelming majority being nonsynonymous (McDonald et al., 2016). We observed that only 12% of mutations reached fixation, and of those fixed mutations, 91% were nonsynonymous. This is suggestive of recombination, however, further work is required to investigate the contribution of genetic exchange during niche- breadth evolution (e.g., analysis of linkage disequilibrium). For example, time-sampled sequencing and analysis of linkage disequilibrium would provide a better understanding of the dynamics of molecular evolution, along with a quantitative assessment of recombination during niche-breadth evolution.

**References**

Bartual, S. G., Otero, J. M., Garcia-Doval, C., Llamas-Saiz, A. L., Kahn, R., Fox, G. C., & van Raaij, M. J. (2010). Structure of the bacteriophage T4 long tail fiber receptor-binding tip. *Proc Natl Acad Sci U S A, 107*(47), 20287-20292. doi:10.1073/pnas.1011218107

Bedhomme, S., Lafforgue, G., & Elena, S. F. (2012). Multihost experimental evolution of a plant RNA virus reveals local adaptation and host-specific mutations. *Mol Biol Evol, 29*(5), 1481-1492. doi:10.1093/molbev/msr314

Carrasco, P., de la Iglesia, F., & Elena, S. F. (2007). Distribution of fitness and virulence effects caused by single-nucleotide substitutions in Tobacco Etch virus. *J Virol, 81*(23), 12979-12984. doi:10.1128/JVI.00524-07

Coleman, J. R., Papamichail, D., Skiena, S., Futcher, B., Wimmer, E., & Mueller, S. (2008). Virus attenuation by genome-scale changes in codon pair bias. *Science, 320*(5884), 1784-1787.

Cuevas, J. M., Domingo-Calap, P., & Sanjuan, R. (2012). The fitness effects of synonymous mutations in DNA and RNA viruses. *Mol Biol Evol, 29*(1), 17-20. doi:10.1093/molbev/msr179

Fokine, A., Chipman, P. R., Leiman, P. G., Mesyanzhinov, V. V., Rao, V. B., & Rossmann, M. G. (2004). Molecular architecture of the prolate head of bacteriophage T4. *Proceedings of the National Academy of Sciences of the United States of America, 101*(16), 6003-6008.

Fokine, A., Zhang, Z., Kanamaru, S., Bowman, V. D., Aksyuk, A. A., Arisaka, F., . . . Rossmann, M. G. (2013). The molecular architecture of the bacteriophage T4 neck. *J Mol Biol, 425*(10), 1731-1744. doi:10.1016/j.jmb.2013.02.012

Ishii, T., & Yanagida, M. (1977). The two dispensable structural proteins (soc and hoc) of the T4 phage capsid; their purification and properties, isolation and characterization of the defective mutants, and their binding with the defective heads in vitro. *Journal of molecular biology, 109*(4), 487-514.

Kostyuchenko, V. A., Leiman, P. G., Chipman, P. R., Kanamaru, S., van Raaij, M. J., Arisaka, F., . . . Rossmann, M. G. (2003). Three-dimensional structure of bacteriophage T4 baseplate. *Nat Struct Biol, 10*(9), 688-693. doi:10.1038/nsb970

Kostyuchenko, V. A., Navruzbekov, G. A., Kurochkina, L. P., Strelkov, S. V., Mesyanzhinov, V. V., & Rossmann, M. G. (1999). The structure of bacteriophage T4 gene product 9: the trigger for tail contraction. *Structure, 7*(10), 1213-1222.

Leiman, P. G., Arisaka, F., van Raaij, M. J., Kostyuchenko, V. A., Aksyuk, A. A., Kanamaru, S., & Rossmann, M. G. (2010). Morphogenesis of the T4 tail and tail fibers. *Virol J, 7*(355), 1-28.

Letarov, A., Manival, X., Desplats, C., & Krisch, H. M. (2005). gpwac of the T4-type bacteriophages: structure, function, and evolution of a segmented coiled-coil protein that controls viral infectivity. *Journal of bacteriology, 187*(3), 1055-1066. doi:10.1128/JB.187.3.1055-1066.2005

McDonald, M. J., Rice, D. P., & Desai, M. M. (2016). Sex speeds adaptation by altering the dynamics of molecular evolution. *Nature, 531*(7593), 233-236. doi:10.1038/nature17143

Ren, Z.-j., & Black, L. W. (1998). Phage T4 SOC and HOC display of biologically active, full-length proteins on the viral capsid. *Gene, 215*(2), 439-444.

Stoltzfus, A., & McCandlish, D. M. (2017). Mutational Biases Influence Parallel Adaptation. *Mol Biol Evol, 34*(9), 2163-2172. doi:10.1093/molbev/msx180

Terzaghi, B. E., Terzaghi, E., & Coombs, D. (1979). The role of the collar/whisker complex in bacteriophage T4D tail fiber attachment. *Journal of molecular biology, 127*(1), 1-14.

Tétart, F., Repoila, F., Monod, C., & Krisch, H. (1996). Bacteriophage T4 host range is expanded by duplications of a small domain of the tail fiber adhesin. In: Elsevier.

Wood, W. B., Eiserling, F. A., & Crowther, R. A. (1994). Long tail fibers: genes, proteins, structure and assembly. In J. D. Karam (Ed.), *Molecular Biology of Bacteriophage T4*. Washington, D. C. : American Society for Microbiolgy.

Yap, M. L., Klose, T., Arisaka, F., Speir, J. A., Veesler, D., Fokine, A., & Rossmann, M. G. (2016). Role of bacteriophage T4 baseplate in regulating assembly and

infection. *Proc Natl Acad Sci U S A, 113*(10), 2654-2659. doi:10.1073/pnas.1601654113
Yu, F., & Mizushima, S. (1982). Roles of lipopolysaccharide and outer membrane protein OmpC of Escherichia coli K-12 in the receptor function for bacteriophage T4. *Journal of bacteriology, 151*(2), 718-722.
